# Isolation and characterization of live yeast cells from ancient vessels as a tool in bio-archeology

**DOI:** 10.1101/549840

**Authors:** Tzemach Aouizerat, Itai Gutman, Yitzhak Paz, Aren M. Maeir, Yuval Gadot, Daniel Gelman, Amir Szitenberg, Elyashiv Drori, Ania Pinkus, Miriam Schoemann, Rachel Kaplan, Tziona Ben-Gedalya, Shunit Coppenhagen-Glazer, Eli Reich, Amijai Saragovi, Oded Lipschits, Michael Klutstein, Ronen Hazan

## Abstract

Ancient fermented food has been studied based on recipes, residue analysis and ancient-DNA techniques and reconstructed using modern domesticated yeast. Here, we present a novel approach. We hypothesize that enriched yeast populations in fermented beverages could have become the dominant species in storage vessels and the descendants of these yeast could be isolated and studied long after. To this end, using a pipeline of yeast isolation from clay vessels developed here, we screened for yeast cells in beverage-related and non-related ancient vessels and sediments, from several archeological sites. We found that yeast cells could be successfully isolated specifically from clay containers of fermented beverages. Genomic analysis revealed that these yeast are similar to those found in traditional African beverages. Phenotypically, they grow similar to modern-beer producing yeast. Both strongly suggesting that they are descendants of the original fermenting yeast. These findings provide modern microorganisms as a new tool in bio-archeology.

**Importance:** So far, most of the study of ancient organisms was based mainly on the analysis of ancient DNA. Here we show that it is possible to isolate and study microorganisms, yeast in this case, from thousands of years old clay vessels, used for fermentation. We demonstrate that it is highly likely that these cells are descendants of the original yeast strains which participated in the fermentation process and were absorbed into the pottery vessels. Moreover, we characterize the isolated yeast their genome and the beer they produce. These results open new and exciting avenues in the study of domesticated microorganisms and contribute significantly to the fields of bio and experimental –archeology that aims to reconstruct ancient artifacts and products.

## Introduction

Experimental archaeology is a field of research that studies ancient cultures by trying to reconstruct ancient lifestyle including tools, housing, cloths and diet (9, 36). Among the most challenging subjects of study in this field are fermented food products such as cheese and pickles, and alcoholic beverages including wine, beer and mead (honey wine). All these products played important roles in ancient societies (46), as a central component of ancient diets, especially important due to their preservation under diverse conditions. In particular, alcoholic beverages filled various important social, political, economic, and religious functions (57). In fact, alcohol, has served throughout history, and continues today, to be an important “social lubricant” in diverse human social and political contexts (16, 33, 39, 65, 74). There is sundry archaeological evidence of fermented beverages, as well as their production and consumption, in ancient societies throughout the world, from late Prehistoric periods onwards (37). Extensive evidence of wine and beer production in Egypt, Mesopotamia and the Near East as early as the mid-4th millennium BCE has been discovered (1, 27, 57). This includes textual evidence in the form of administrative lists and narratives that mention such beverages, including actual recipes of different types of wine (43) and beer, as well as small scale models and paintings of their production (15, 57). Similarly, chemical evidence of wine and beer production has been found in the form of various components of breweries (60) including vessels and related installations,. Such residue analyses have enabled the identification of alcoholic beverages of numerous cultures as early as the Neolithic period, (ca. 6,000-5,000 BC) in the region of modern Georgia (56) to ancient China (83, 84), Mediterranean France (7), Cyprus (15), Bronze and Iron Age Israel (43), Nordic cultures of Scandinavia (59) early Celts in Germany (78), early cultures of the Andes (65), Prehistoric Europe and Indo-Iranian Asia (19), and Egypt (73), leading in some cases to the identification of specific compositions of these beverages (37, 58, 74).

Based on this evidence, there have been several attempts to recreate ancient beer and wine, but those were always brewed using modern ingredients combined with modern domesticated commercial yeast (predominantly *Saccharomyces cerevisiae*) (57, 78) and not with the actual microorganisms which might have been used in the production of these fermented beverages. On the other hand, up until now, the study of ancient microorganisms, including bacteria (3, 17), viruses (61) and yeast (13) was mainly focused on ancient DNA studies.

Here, we isolated yeast directly from ancient vessels, which had previously been suggested to have served as beverage containers. We found that yeast are significantly more abundant in these putative beverage containers, than in other non-beverage-related archaeological vessels, from these and other sites, or in sediments from these sites and the surrounding environment. This supports the hypothesis that the yeast found in the beverage containers originated from the large amount of yeast cells that grew during the beverage fermentation and continued to reproduce and survived as colonies in the micro-environments of the pores in the ceramic matrix of these vessels. In agreement with this hypothesis, phenotypic and genomic characterization of these yeast strains, including genomic DNA sequencing, showed that they are similar to yeast found in traditional beers, and are able to ferment and produce drinkable beer similar to modern beverages.

## Results

### Isolation of yeast strains from ancient vessels

We hypothesized that enrichment of clay vessels with large amounts of fermenting yeast which were absorbed into the vessel pores of the ceramic matrix permanently changed the vessel’s microorganism content (vessel microbiome).

Indeed, testing several modern vessels which were filled with filtered and un-filtered beer and buried for three weeks underground, and further tests of a wine clay vessel that was unused for more than two years revealed that yeast cells can be found in the clay matrix, after an extended period of time (Fig 1A). Next, we tested several methods of yeast isolation and developed a pipeline (Fig. 1B) that enabled us to efficiently isolate viable yeast cells from these modern clay containers. In contrast, we could not isolate any live yeast from the control vessels which were filled with filtered beer, nor were yeast cells detected by electron microscopy (Fig. 1A, left panel). Next, we tested ancient ceramic vessels from three different historical periods, found in four different archaeological sites located in Israel (Fig. 2A). Each of these sites contained vessels that were assumed to have been associated with fermented beverages, based on ancient iconography, functional analysis based on the vessels’ shape, or previously conducted organic residue analysis. Electron microscopy visualization showed “yeast like” structures (Fig. 2C) similar to that of the modern clay vessels (Fig. 1A) which prompted us to try isolating live yeast cells from the ancient vessels.

**Fig. 1.**
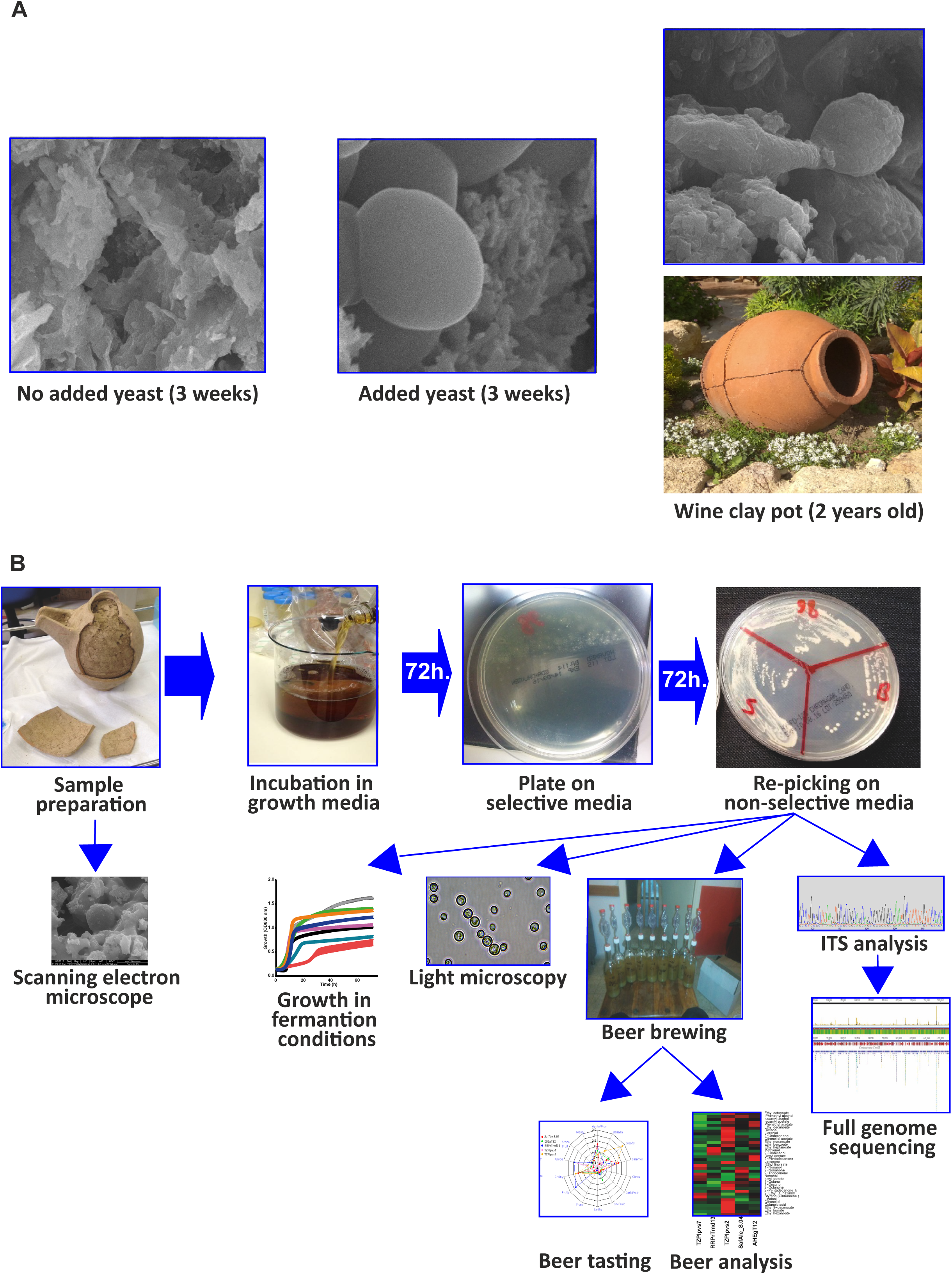
Isolation of yeast from clay vessels. **(A)** Yeast strains in clay vessels. Scanning electron microscope (SEM) pictures of the inside of a modern clay vessel buried in the ground for three weeks without beer (left panel) and pre-soaked with unfiltered beer (middle panel) prior to burial. On the right panel is a two-year-out of use wine clay vessel (bottom) which yielded live yeast cells observed as colonies and by EM (upper). Yeast cells were only successfully isolated from the last two vessels. **(B)** The pipeline of yeast isolation and characterization from vessels. Putative fermented beverage-containing vessels were carefully dismantled. Small pieces were sent for SEM and the rest were incubated in growth medium (YPD) for 72 hours at room temperature. Samples were plated on selective plates with antibiotics to eliminate bacteria. After 72 hours, yeast colonies appeared and were regrown on new plates. The yeast strains were taken for various analyses including full genome sequencing and comparison of growth in fermentation related conditions in beer wort. In addition, beer was brewed according to a standard recipe using the isolated yeast strains. The presence of aromatic and flavor compounds in the beers was analyzed quantitatively and their flavor was qualitatively evaluated by specialized beer tasters.

**Fig. 2.**
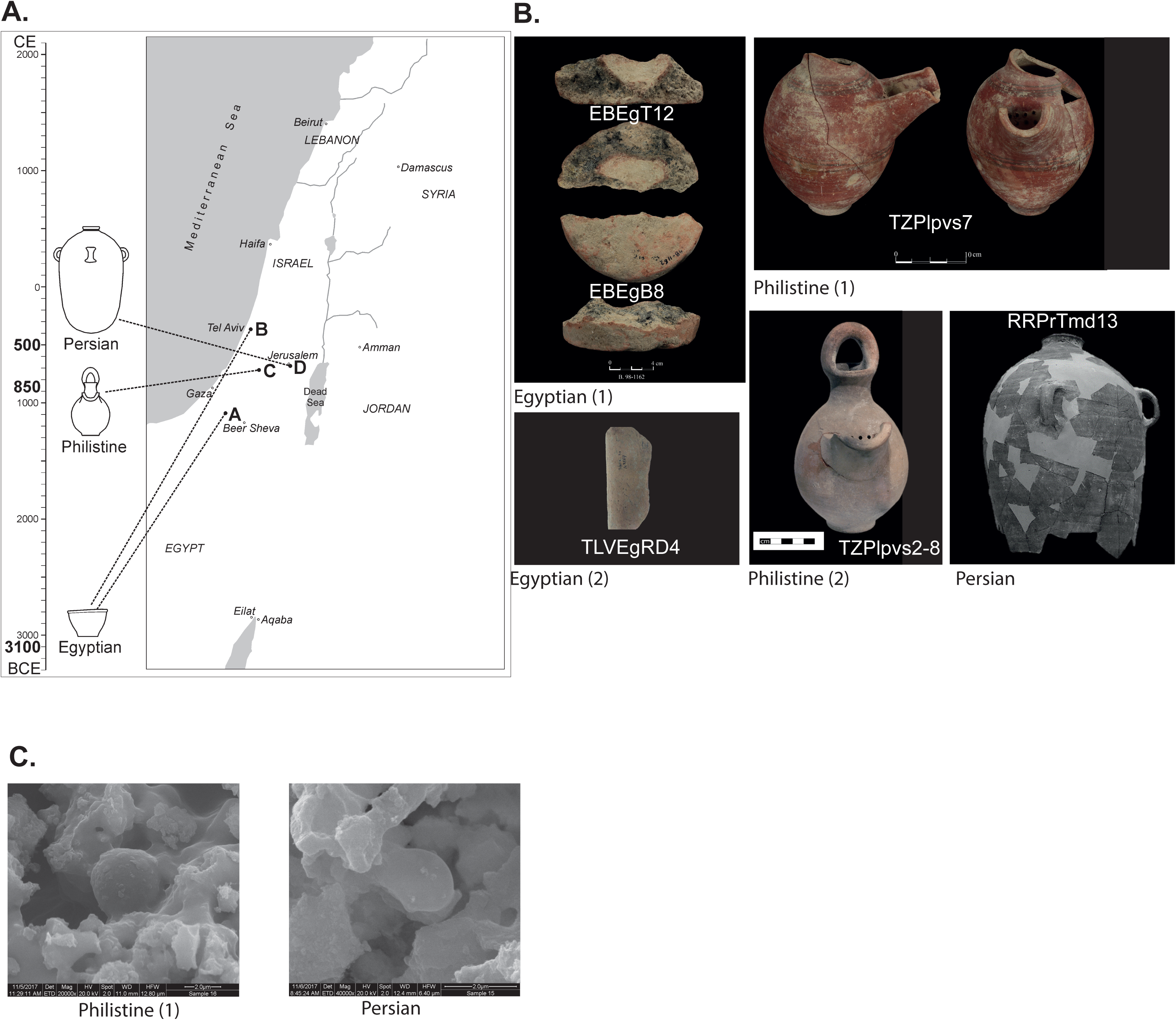
Ancient vessels that putatively contained fermented beverage and were used for yeast isolation. **(A)** A map and timeline of the archaeological sites from which the vessels yielding fermenting yeast strains were excavated. **(B)** Pictures of the vessels. The white text in the pictures indicates the name of the yeast strain isolated and the text below the pictures denotes the archaeological culture that the vessels are associated with. **(C)** Representative SEM picture of vessels with “yeast like” structures (Compare to Fig. 1A).

The first vessels were excavated from two sites dated to the Early Bronze Age IB (ca. 3100 BCE). The first site is En-Besor in the northwestern Negev desert, a site relating to the Egyptian activities in southern Canaan during the late 4th millennium BCE, as evidenced through typical Egyptian architecture, pottery, and clay bullae with hieroglyphic symbols (25, 75). The second site was recently excavated at Ha-Masger Street in Tel Aviv, and contained basin fragments, typical of Egyptian-style breweries, perhaps evidence of an Egyptian enclave within a local Canaanite settlement. We tested five ceramic fragments (Fig. 2A, B) of vessels from these two sites, that according to ancient Egyptian depictions were used as beer basins (57). These vessel fragments yielded three yeast strains, two from En-Besor, and one from Ha-Masger Street, designated EBEgT12, EBEgB8 and TLVEgRD4 (Table 1).

**Table 1.**
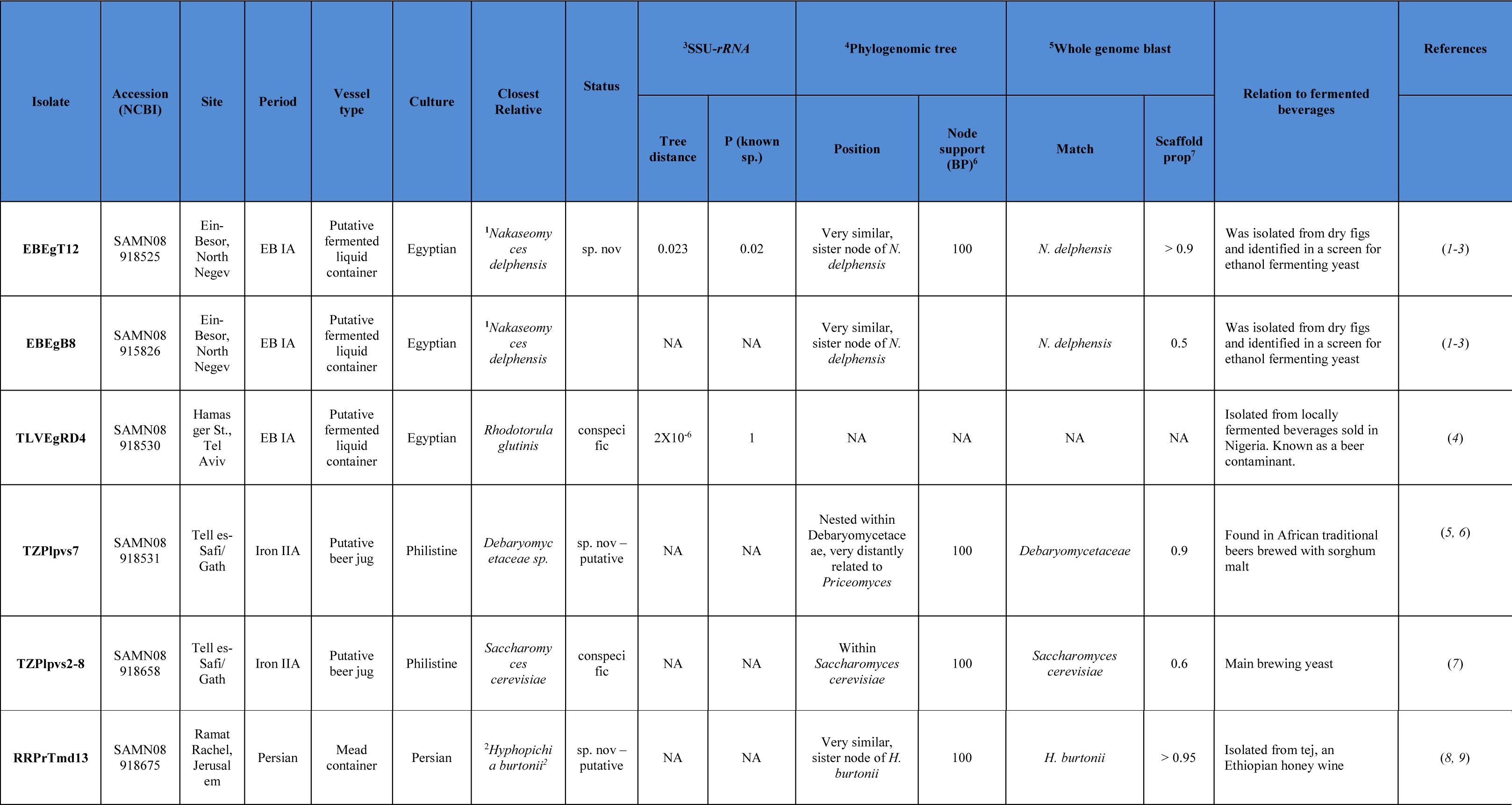
Genetic identification of yeast strains isolated from ancient vessels. Summary of the phylogenetic identification of the yeast strains isolated from ancient putative beverage vessels and the source, site and period of the vessels. SSU-*rRNA*: The patristic distance (substitutions per base) between the isolate and its closest relative, given with the probability that this is an intraspecific tree distance.

The third site sampled was Philistine Tell es-Safi/Gath (in central Israel), specifically from contexts dating to the Iron IIA (ca. 850 BCE) (54, 55, 63). The Philistines, one of the so-called “Sea Peoples,” were an important culture in the Levant during the Iron Age (ca. 1200-600) and are often mentioned in the Bible as enemies of the Israelites (76). At the time, Philistine Gath was the largest and most important Philistine site in the region (54). We tested 12 samples from two well-preserved Philistine jugs (Fig. 2B and SI Appendix, table S1), usually associated with beer or other fermented alcoholic drinks, based on their spout and a strainer-spout on their side (18, 30, 42). Each of these vessels yielded a yeast strain, designated TZPlpvs7 and TZPlpvs2-8 (table 1). The fourth site was Ramat Rachel, located between Jerusalem and Bethlehem (Fig. 2A) (49). During the Iron Age and Persian Periods (ca. 8^th^-4^th^ cent. BCE) it sequentially served as the residence of the local representative of the Assyrian, Babylonian and Persian empires, as a center for tax collection, and for diacritical feasting events (21). From this site we examined four storage jars, typical of the Judean region during the early Persian period (Fig. 2B), all found in a refuse pit, and which contained mead according to previous organic residue analyses (48). One of these pot-shards yielded a yeast strain designated RRPrTmd13.

In summary, we succeeded in isolating six yeast strains from 21 beer- and mead-related ancient vessels (Table 1 and (SI Appendix, Table S1).

### Negative controls

One of the key questions in the current research is whether the yeast cells descended from the enriched ancient yeast cultures which fermented the liquid stored in the excavated vessels, or are they equally abundant in the environment? In order to answer this question, we used the above method to isolate yeast from non-beverage related vessels and sediments from the surrounding environment of the excavated site. To this end, we tested 27 samples from other ancient vessels from these same sites, vessels which were not associated with beverage storage, but with other functions, including cooking pots, petite jugs, lamps and bowls. None of these vessels yielded yeast (SI Appendix, Table S1), save for the lamps (on this, see below). Furthermore, we tested 53 samples of sediments and stones gathered from these archeological sites, adjacent to the locations where the putative beverage vessels were found. These samples yielded two yeast strains, one from a stone from En-Besor (SI Appendix, Table S1, sample 26) which was identified by ITS analysis as the pathogen *Candida albicans* (Table S2) and is presumably a contamination originating from humans. The other one from a sediment sample from Tell es-Safi/Gath (SI Appendix, Table S1, sample 103), has an unidentified ITS and is probably an undescribed wild yeast. Last, since yeast is often associated with plants (51), we also tested 30 samples of sediment and stones from the non-archeological site Ma’on, as well as agricultural fields in the proximity of the sites of Tell es-Safi/Gath and Ramat Rachel. We were unable to isolate any yeast strains from these samples using our pipeline (SI Appendix, Table S1). Overall, we found two yeast strains out of 110 non-beverage related control samples. Thus, the findings of six yeast strains from 21 samples of putative fermented beverage vessels versus two yeast strains from 110 control samples is significant and hardly incidental (with Fisher’s exact test *P-value* of 0.0006).

### Genome sequencing of the isolated yeast

In addition to the ITS region identification carried out on all isolated yeast (Table S2), we sequenced the full genomes of the six yeast strains that had been isolated from beverage associated ancient vessels (For accession numbers see Table 1). We also sequenced the genome of one of the yeast strains that was isolated from the controls, RRPrNerP7 (accession: SAMN08918674), which was isolated from an oil lamp found at Ramat-Rachel (SI Appendix, Fig. S1). All yeast strains were identified based on similarities to yeast strain genomes from the NCBI database (SI Appendix, Fig. S2) and there was a match between the ITS identification and the full genome sequencing.

The two yeast strains EBEgT12 and EBEgB8, which were isolated from the Egyptian vessels excavated at En-Besor, are genetically close to one another, and show high similarities to *Nakaseomyces delphensis* (also known as *Saccharomyces delphensis*) (Fig. 3A and Table 1), which was isolated from dry African figs and is not common in soil (81). This supports the notion that the yeast cells originated from the vessels themselves, and not the environment, and suggests that perhaps figs were used in the fermented beverage production. Additionally, based on LSU-*rRNA* barcoding analysis, these strains appear to belong to an unrecorded species (supplementary material and Table 1). To draw functional insight from the genomes of EBEgT12 and EBEgB8, we identified 596 orthologous gene clusters with copy number variation between the two isolates. Of these genes, we further compared 79 orthologous gene clusters of genes that were related to transmembrane transport and metabolism of various carbohydrates and were previously described as having copy number variations in beer producing yeast strains (23). Despite the overall high genetic similarities between these two yeast strains (Fig. 3A), EBEgT12 had 67 genes with the expected duplications or deletions characteristic to beer yeast strains (SI Appendix, Table S4), whereas only 12 occurred in while EBEgB8, which did not produce drinkable beer on its own (see below). This data suggests that EBEgT12 was better adapted for beer production than EBEgB8, as indeed was observed while producing beer from these yeast strains (see below, beer production section).

**Fig. 3.**
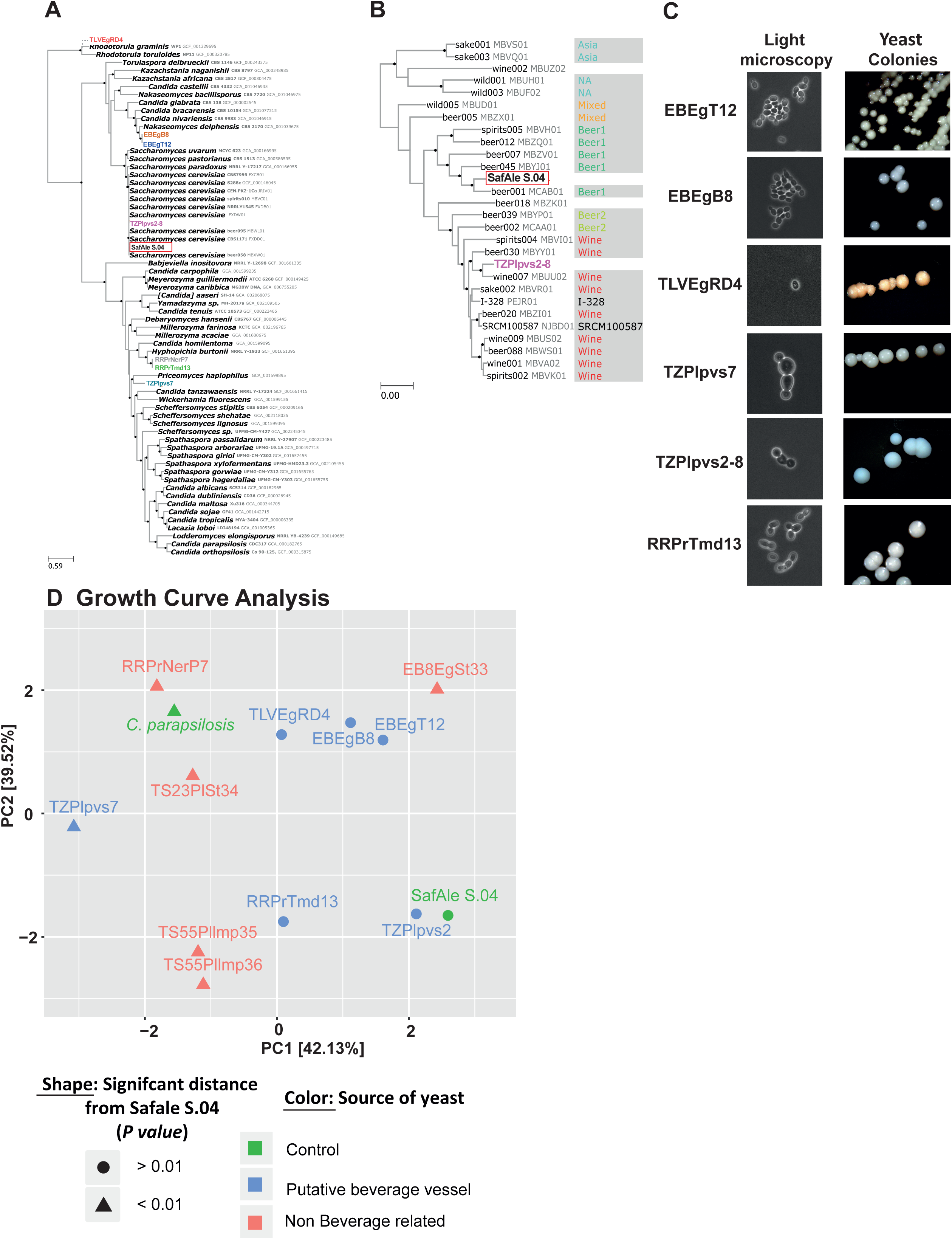
Genotypic and phenotypic characterization of yeast strains isolated from ancient vessels. **(A-B)** Phylogenetic trees, based on full genome sequencing of the isolated yeast strains. Black bullets at nodes represent maximal bootstrap percentage node support. The newly isolated strains are in color font. The modern beer yeast species *Saccharomyces cerevisiae* (SafAle S.04), which served as control, is surrounded by a red square. (**A**) 118 gene partitions and a representation of *Saccharomycetaceae + Debaryomycetaceae*. (**B**) Comparison of TZPlpvs2-8 (in purple) to modern wine and beer strains, based on 465 gene representatives and partitioning of *Saccharomyces cerevisiae*. Reference strains are denoted by NCBI strain name and accession number followed by clade affiliation (23). (**C**) The shape of the isolated yeast cells in light-microscopy (left panel) and colonies on YPD agar plates (right panel). **(D) Growth Curves analysis.** The yeast strain isolated from putative beverage vessels grow in beer wort with similar kinetics to a modern beer-yeast. Principal component analysis (PCA) of the distances of the growth curves of yeast grown in beer wort under fermentation related conditions (SI Appendix, Fig. S3). The modern domesticated beer yeast strain SafAle S.04 served as positive control and the pathogenic yeast *C. parapsilosis*, served as the negative control. The marker’s shape denotes the statistical significance of the distance from SafAle S.04 growth curve kinetics and the color denotes the source of the yeast: control, putative beverage container or non-beverage-related vessel.

TLVEgRD4, the 3^rd^ yeast strain which was isolated from the Egyptian vessels, showed high similarity (Fig. 3A and Table 1) to the red pigmented yeast *Rhodotorula glutinis* which is a known food contaminating agent found in Nigerian and other beers (20, 38).

From one of the Philistine vessels, we isolated yeast strain TZPlpvs7, which was found to be similar to yeast strains of the *Debaryomycetaceae* family (Fig. 3A and Table 1). Members of this family were isolated from traditional African beers brewed with sorghum malt (31, 53). The second yeast strain isolated from the other Philistine vessel, TZPlpvs2, is *Saccharomyces cerevisiae,* which is the most commonly used species of domesticated yeast, and plays a central role in modern beer, wine and bread industries (47). We further tested the similarity of TZPlpvs2 to known beer and wine producing *S. cerevisiae* strains, and found that it is close to strain Wine-007 (NCBI assembly MBUU02, Fig. 3B), a modern yeast strain used in wine production (23).

RRPrTmd13 which was isolated from a mead containing vessel (based on organic residue analysis), was found to be similar to the yeast *Hyphopichia burtonii* (*Endomycopsis burtonii*) (Fig. 3A and Table 1). Significantly, this yeast species was previously isolated (5, 19) from *tej,* an Ethiopian honey wine (2) – a type of traditional African mead.

We also sequenced the genome of the yeast strain RRPrNerP7 (NCBI Accession: SAMN08918674) that was isolated from a clay oil lamp (SI Appendix, Fig. S1) from Ramat-Rachel. Surprisingly, its sequence was found to be similar to *H. burtonii* (Fig. 3A) like RRPrTmd13, the mead vessel yeast strain which was isolated from the same site. Nevertheless, RRPrNerP7 and RRPrTmd13 were divergent from each other in phenotypes related to several beverage production aspects. We compared RRPrTmd13 and RRPrNerP7 regarding duplications and deletions in 52 orthology clusters with gene ontologies related to the metabolism of various carbohydrates and the transmembrane transport of iron, sodium, and sugars, found to have characteristic copy number variations in modern wine producing *S. cerevisiae* yeast (23), although not specifically studied in mead producing yeast strains. As can be seen in SI Appendix, Table S4, the duplications and deletions that occurred in both isolates are significantly different (ttest, *p-value* = 2×10^−7^). Additional phenotypic differences between RRPrTmd13 and RRPrNerP7 are described below.

### Genome wide BLASTn

Each genome assembly was analyzed with the online version of BLASTn. To summarize the results, we considered all the taxonomic IDs that constituted more than 5% of the matches (Table 1, Fig S2). For Isolate TZPlpvs7, over 50% of the scaffolds matched to one of three *Debaryomycetacid* species with only 80% identity (stdev 4.1%) to all the species, and over 40% additional scaffolds matched *Debaryomycetacid* species with lower identities. Similarly, the best match of almost all the scaffolds of either isolate RRPrTmd13 and RRPrNerP7 was *H. burtonii*, albeit with a mean percent identity of only 84.5% (stdev 4.5%). Finally, the best matches of 50% and 97% of the scaffolds of isolates EBEgB8 and EBEgT12, respectively, were in the *Nakaseomyces*/*Candida* clade, but with a low mean percent identity of only 83% (stdev 6%). We would thus suggest that these five isolates represent species that are not yet recorded in the NCBI nucleotide repository. Conversely, for TZPlpvs2-8, over 60% of the scaffolds had 99.9% identity (stdev 0.14%) with *S. cerevisiae*, indicating that this isolate is very similar to records of S. *cerevisiae*, in agreement with the phylogenomic analysis.

### Phylogenomic analysis

To validate the phylogenetic position of the isolates, we selected reference genome assemblies of 55 isolates that are available on Genbank (see Fig 3a for accession numbers). We then annotated coding sequences in the reference genomes as well as in our isolate genome assemblies, using Augustus 3.2.3 (41). For the annotation process, we have chosen the coding sequences of the nearest available reference relative as hints (see Fig. 3a), and either *Saccharomyces* or *Candida tropicalis* as the model species for *Saccharomycetaceae* and *Debaryomycetaceae* species respectively. We extracted a protein sequence file for each isolate genome and reference genome and assigned orthology information to each gene with eggNOG 4.5.1 (35) (http://eggnogdb.embl.de/#/app/home). We selected orthologs with one representative in at least 50% of the reference genomes and in at least three out of our five isolates. Protein sequences of each ortholog were aligned with MAFFT (40) using the L-ins-i algorithm and each ortholog alignment was trimmed with TrimAl using the gappyout algorithm. Using treeCl (26), we reconstructed maximum likelihood gene trees for each ortholog, and clustered the resulting gene trees based on the Weighted Robinson Folds (WRF) (69) pairwise inter-tree distances and the db-scan clustering algorithm, to assess the existence of conflicting phylogenetic signals. For every cluster, treeCl produces a supermatrix of all the genes in the cluster, which we used for a partitioned tree reconstruction with RAxML (77) using the LG evolutionary model and 100 thorough bootstrap replicates for branch support *Saccharomyces cerevisiae*. To recover the phylogenetic position of two *S. cervisiae* isolates (Sefale-S04 and TZPlpvs2-8), we repeated the workflow described above, using a targeted reference dataset of 26 *S. cervisiae* genomes covering the diversity of known isolates, as described in Gallone et al. (2016). In this case we retained one to one orthologs represented in at least 70% of the reference genomes and in both our isolates. Due to the high sequence identity among the analyzed genomes, all belonging to *S. cervisiae*, we retained only the most informative 650 orthologs by selecting alignments with at least 10 unique sequences and at least 10 parsimony informative alignment columns (i.e., at least two character states in the column, each occurring in at least two sequences).

The sequence alignments of 118 orthologs passed our filters and were included in the analysis of the *Saccharomycetaceae* + *Debaryomycetaceae* dataset. Conflicting phylogenetic signals were not detected among them, as the db-scan algorithm has detected only one cluster, which was robust to changes in minimal local radius cutoff. The phylogenetic tree was reconstructed from a supermatrix of all the 118 orthologs (Fig 3a) with the matrix and partition (uploaded files 6 and 7). Isolates EBEgT12 and EBEgB8 were very similar with less than 4×10^−4^ substitutions per base (SPB). They clustered as the sister clade of *Nakaseomyces delphensis*, but with much larger sequence divergence (over 0.13 SPB). Isolate TZPlpvs7 was resolved as a Debaryomycetaceae sp., which is divergent from other con-familials for which a genome assembly is available (at least 0.8 SPB). Isolates RRPrTmd13 and RRPrNerP7 were very closely related to each other (less than 4×10^−4^ SPB) and emerged as a sister clade of *Hyphopichia burtonii* (Debaryomycetaceae) with a sequence divergence of over 0.2 SPB. Isolate TZPlpvs2-8 clustered within the *S. cerevisiae* clade (Fig 3a) with maximal node support. Based on our *S. cerevisiae* focused phylogenomic analysis (Fig 3b) TZPlpvs2-8 is a part of the Wine cluster as was recovered by (23). This cluster originally included both beer and wine yeasts. It is most closely related to isolate “Wine007” with maximal node support, and with sequence divergence of 8×10^−4^ SPB. This sequence divergence is larger than those observed between RRPrTmd13 and RRPrNerP7 or between EBEgT12 and EBEgB8 and is not contrary to observed phenotypic differences. The *S. cerevisiae* focused analysis included 650 orthology clusters that passed the filtering steps (see Materials and Methods section). In this analysis, the number of gene tree clusters was maximized when using a minimal local radius of 0.03 in the db-scan analysis, and resulting in two tree groups of 465 and 185 trees. Fig 3b is based on the larger group, whereas the smaller group yielded a tree with a similar topology, but with an overall shorter tree distance. It is thus an artifact of the weighting procedure of the WRF parameter and does not represent a real phylogenetic conflict. As we observed different phenotypes in isolates EBEgT12 and EBEgB8, with only isolate EBEgT12 producing beer, we expected that this difference will be reflected in gene copy number variation (CNV) between the beer producing yeast and the non-producing yeast, as previously shown by Gallone *et al.* (23). Despite the overall high genetic similarities between these two yeast strains, we identified 79 orthology clusters with CNV between the two isolates, which were related to transmembrane transport and metabolism of various carbohydrates, also described by (23) as having CNV in beer producing yeasts. EBEgT12 had 67 genes with the expected duplications or deletions in beer yeasts (SI Appendix, Table S4), while EBEgB8 had only 12, supporting EBEgT12 as better adapted for beer production than EBEgB8.

Additionally, we observed different phenotypes in isolates RRPrTmd13 and RRPrNerP7, with only isolate RRPrTmd13 producing mead. In this case we also expected that this difference will be reflected by CNV instances between the two isolates. Although Gallone et al. (2016) did not analyze mead producing yeasts, wine shares some of the sugar sources with honey based mead, and similarly, the mead producing isolate (RRPrTmd13) shares some of the duplications and deletions with the wine producing isolate (23) when compared with isolate RRPrNerP7 (SI Appendix, Table S1). In this case, however, both isolates had similar numbers of CNV instances expected in wine yeasts (27 and 25 for RRPrTmd13 and RRPrNerP7 respectively), providing no prediction as to the expected phenotype.

### Taxonomic identity of isolates based on LSU-*rRNA* barcoding

Taxonomic classification of isolates EBEgT12 and TLVEgRD4 was assessed *via* LSU-*rRNA* barcoding, as this marker has been comprehensively sampled across the taxonomy of Ascomycota, and is more variable than the SSU-*rRNA* gene (68). BLASTn 2.6.0+ (10) was used to identify the LSU-rRNA locus in each genome assembly, with the LSU-*rRNA* SILVA (68) database sequences as query and the genome assemblies as target. In each genome assembly, the best match to any of the target sequences in each assembly was recovered as the isolate’s LSU-*<u>rRNA</u>* gene. We further composed a relevant reference dataset by running a second BLASTn analysis, in which the isolate LSU-*rRNA* sequences were used as queries, and the LSU-*rRNA* SILVA database as target. The best 500 matches to each of the isolate sequences were retained, and redundancies were eliminated by retaining only the centroid sequences of 99% identical clusters, as predicted with VSEARCH v2.4.3 (71). The resulting dataset, together with the isolate LSU-*rRNA* sequences recovered from the genome assemblies, was used in a phylogenetic analysis to identify the phylogenetic position of the isolates. The sequences were aligned with MAFFT v7.310 (40), positions with over 0.8 gap proportion were removed with TrimAl v1.4.rev15 (11), and a phylogenetic tree was built with RAxML 8.2.10 (77), using the GTRGAMMA model and 100 replicates of rapid bootstrap trees for node support.

To check whether the isolates belonged to established species we calculated all the intra-species patristic distances (the cumulative branch length between two tree nodes) in the LSU-*rRNA* phylogenetic tree and computed their distribution. We then calculated the patristic distance between each isolate for which LSU-*rRNA* was recovered and its closest relative and tested whether this distance belonged to the distribution of the intraspecific patristic distances. Patristic distances were computed with ETE 3 (34). Our trimmed non-redundant LSU-*rRNA* sequence alignment included 350 reference sequences and the isolates EBEgT12, and TLVEgRD4 (Table 1), with 853 positions and less than 0.1% missing data. The redundant dataset, as well as the non-redundant alignment and the trimmed alignment are included (uploaded files 1-4). The maximum intraspecific patristic distance in our resulting tree was 0.012 substitutions per base (SPB). Only isolate EBEgT12, was divergent enough from its closest relative (*Nakaseomyces delphensis*; *Saccharomycetaceae*) to constitute a novel species (0.023, *P-value* = 0.02). It is worth to note that by removing redundant sequences we over-estimated the *P-value* and this result is thus very conservative. Isolate TLVEgRD4 was found to be identical to *Rhodotorula glutinis* (*Sporidiobolaceae*, 2×10^−6^ SPB). The LSU-*rRNA* phylogenetic tree is in uploaded file 5. The results of all the analyses are summarized in Table 1.

### Phenotypic characterization of the isolated yeast

We compared several phenotypes of the isolated yeast strains related to alcoholic beverage production. As a positive control we used the modern, commercially available beer yeast strain, *S. cerevisiae* SafAle S.04 (Fermentis Division of S.I.Lesaffre, France). First we compared the morphology of cells and colonies (Fig. 3C, left panel), using phase light microscopy to image colonies (Fig. 3C, right panel) on agar plates containing the lab standard yeast medium YPD. All yeast strains showed the common structure of budding yeast cells and white smooth colonies, with the exception of TLVEgRD4 which yielded red colonies. The red pigmentation is in agreement with its identification as *R. glutinis,* which produces several carotenes including β-carotene (28).

Next, we hypothesized that the isolated yeast strains were naturally selected to grow in beverage fermentation conditions and would be able to grow in beer wort, similar to modern domesticated beer yeast strains. To test this hypothesis, we compared growth kinetics of ancient isolated yeast strains to those of the modern beer-yeast-strain SafAle S.04 when grown in wort (SI Appendix, Fig. S3A). As a negative control, we used the pathogenic yeast species *Candida parapsilosis* (79) which, unquestionably, is not used for beverage production. To compare the growth curves, we fit each curve to a logistic equation (SI Appendix, Fig. S3B) which models growth curves (80). Next, we calculated the relative distance between the various fitted equations using principal component analysis (PCA) demonstrating the relative similarities between the parameters of the fitted curves (Fig. 3D). We found that a high correlation (r = 0.95) exists between the growth curve shape and whether the yeast strain was isolated from a putative beer vessel or not. All yeast strains isolated from vessels, which were believed to have originally contained fermented beverages, grew similarly to SafAle S.04, except for TZPlpvs7, while all the other yeast strains, from lamps, sediments and stones, showed different growth kinetics than SafAle S.04 (Fig. 3D). These results suggest that indeed the yeast strains isolated from the putative beverage containers are progenies of yeast which were selected in the past for growth in fermentation related conditions.

### Analysis of beer produced by the isolated yeast

Finally, we tested the ability of the isolated yeast strains to produce drinkable alcoholic beverages. To this end, we performed an initial screen using a standard common recipe of beer brewing (44) with each one of the isolated yeast strains. Strains EBEgT12, TZPlpvs7, TZPlpvs2 and RRPrTmd13 produced aromatic and flavorful beer and were taken for additional compounds and flavor analysis. In contrast, the following yeast strains were excluded from further analysis; EBEgB8, the stone originating yeasts EB8EgSt33 and TS23PlSt34, and the yeasts isolated from the oil lamps RRPrNerP7, TS55Pllmp35 and TS55Pllmp36, which produced beer with mild or strong spoiled aroma and flavors (85). TLVEgRD4 was also excluded, as *Rhodotorula glutinis* was reported to be a pathogenic beer spoiler yeast species (86).

Next, we compared the beer produced by the yeast, which passed the initial screening, to that produced by the positive control, to SafAle S.04 (Fermentis Division of S.I.Lesaffre, France). Comparison of the total carbohydrates (Fig. 4A) and alcohol (Fig. 4B) concentrations produced by yeast showed that besides TZPlpvs7, all the other yeast strains exploit carbohydrates and produced about 6% of alcohol, similar to the “professional” beer yeast strain SafAle S.04. We performed further qualitative analyses of several aromatic and flavor compounds in the various beers by HS-SPME-GCMS (SI Appendix, Table S6). The detected compounds were either known to be present in beers (8, 70, 72) or are other members of the alcohol, ester, monoterpenoid, and carboxylic-acid groups. This analysis shows that relatively high ratios of many aroma compounds were detected in the beer produced using the TZPlpvs2 strain which was no surprise, as this strain was identified as *S. cerevisiae*. Moreover, comprehensive analysis of aromatic and flavor compounds (Table SI Appendix, S6) shows that TZPlpvs2 and EBEgT12, produced beers that clustered with SafAle S.04 (Fig. 4C).

**Fig. 4.**
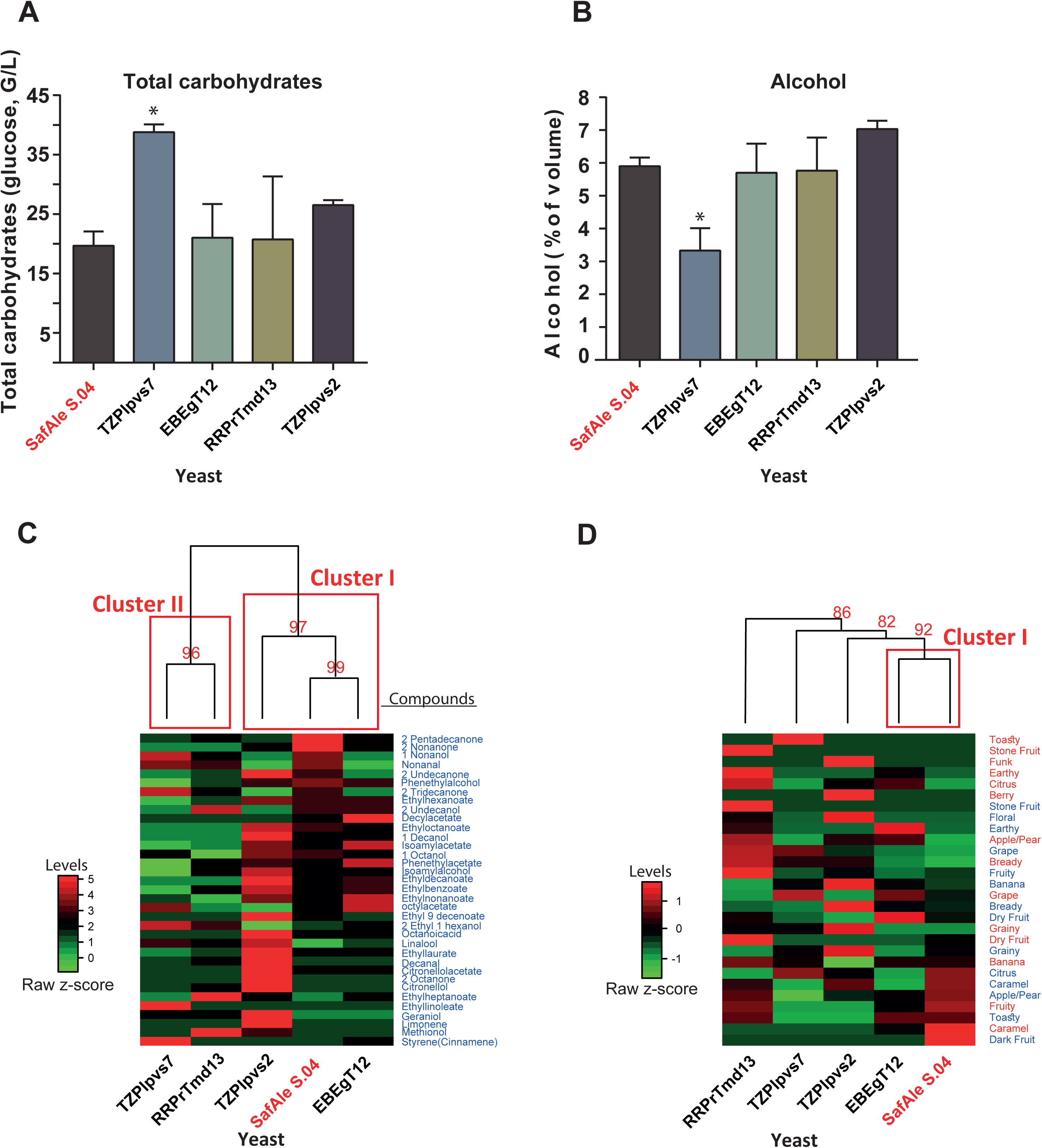
Characterization of reconstructed beer produced by yeast strains isolated from ancient vessels. Beer was brewed using the yeast strains isolated using a standard brewing recipe. The modern beer yeast strain *S. cerevisiae* (SafAle S.04) served as a control**. (A)** Levels of total carbohydrates in the beers (as glucose); **(B)** Amount of alcohol produced; **(C)** Heat map of clustered levels of aromatic and flavor compounds found in the beer. The levels of various compounds were normalized as a percentage of the highest value for each compound; **(D)** Heat map of clustered samples based on parameters of beer tasting of aromas (red text) and flavors (blue text) see also SI Appendix, Fig. S4. In both **C** and **D**, the clustering was performed using Ward’s method with Euclidean distances. In the red text are the approximated unbiased (AU(*P-values* in percentages of the nodes. The red squares denote clusters with *P-value* > 95%

The beers that passed the initial screen were also compared by organoleptic descriptive analysis performed by members of the Beer Judge Certification Program (BJCP, https://www.bjcp.org) beer taster’s organization. The results of the test were in agreement with the chemical analysis: RRPrTmd13, and to a lesser extent, TZPlpvs2, produced beers which are similar in color, aroma and flavor to that of SafAle S.04 (Figs. 4D and SI Appendix, S3).

### Ancient Lamps

An exception to the notion that yeast are significantly isolated from ancient beverage containers in comparison to other vessels were clay lamps, which usually contained olive oil, from both Tell es-Safi/Gath and Ramat-Rachel. Surprisingly, we succeeded in isolating three yeast strains from 6 ancient lamps (SI Appendix, Fig. S1A-C). A possible explanation to the presence of yeast in these lamps might be that the yeast derives from yeast cells existing on olives. These yeast cells were not killed during the cold-press extraction of the oil and they were absorbed into the pores of the clay lamps. This is in agreement with previous observations showing that olive oil contains live yeast cells (14). We also confirmed that yeast cells are indeed present in olive oil by isolating live yeast cells from a modern bottle of olive oil that had been sealed for two years (SI Appendix, Fig. S1D). Finally, we identified the yeast strains by amplification and sequencing of their ITS region and found that the two yeast isolated from the vessels from Tell es-Safi/Gath are strains of *Yarrowia lipolytica* (SI Appendix, Table S2), a yeast strongly associated with oil flora (12, 64) which is not used for beer production (24). Thus, we suggest that the oil lamp results support the notion that yeast colonies remain alive in clay vessels and it is feasible to isolate them.

## Discussion

In this study, we isolated yeast cells from ancient vessels excavated at archaeological sites in Israel. These vessels belong to vessels types that, based on their shape, or, in the case of the vessel from Ramat Rachel, based on organic residue analysis, were considered to have contained fermented beverages such as beer and mead (honey wine).

The main challenge of this research lies in the question whether the isolated yeast strains originated from the ancient yeast which fermented ancient beverages in the archeological vessels, and the yeast cells that were discovered are in fact descendants of the original yeasts, having survived and continued to grow in micro-environments in pores within the clay matrix of the vessels. Or, perhaps, they are wild yeast from the environment or simply a recent contamination. Several lines of evidence strongly suggest that the yeast strains we isolated here are indeed descendants of fermenting yeasts which were enriched in the ancient vessels:

First, the number of isolated yeast strains from putative beverage vessels (six out of twenty-one samples), in comparison with the yeast strains isolated from the control samples (two out of 110 samples), is significantly biased towards the beverage-related vessels;

Second, all yeast strains isolated from the putative beverage vessels, besides TZPlpvs7, grew in beer wort medium, similar to the modern domesticated beer yeast SafAle S.04; while all the yeast strains isolated from the control samples, show different growth parameters under these conditions (Fig. 3);

Third, most of the yeast strains isolated from putative beverage-containers produced drinkable aromatic and flavored beer (Figure 4), while all the control isolated yeast strains produced spoiled aromas and flavors. Thus, there is a strong correlation between the source of isolation (putative beverage vessel or not), similar growth in wort compared to modern beer yeast strains, and the ability to produce drinkable beer. Such a correlation strongly suggests that the yeast strains isolated from putative beverage vessels are descendants of yeast strains that have experienced the selection pressures of alcoholic fermentation and beverage production environment, and continued to reproduce over the ages in micro-environments within the ceramic matrices of the ancient beverage vessels.

Fourth, the molecular phylogeny of the yeast strains isolated from the ancient vessels also supports the notion that they originated from ancient, fermented–liquid-related yeast strains. TZPlpvs2, being *S. cerevisiae*, the major fermenting yeast species today (66), is often found on fruits and flowers and less in soil (50). EBEgT12, EBEgB8 and TZPlpvs7 are similar to yeast species found in various traditional beverages in Africa (31, 53). Yeast strain RRPrTmd13 which was isolated from honey-wine vessel, identified as such through organic residue analysis, is highly similar to a yeast species found in the Ethiopian honey-wine tej (5). Lastly, the two yeast strains isolated from Philistine oil lamps were found to be distinct strains of *Yarrowia lipolytica*, a yeast which tends to grow in olive oil (12, 64). In contrast, the only two yeast strains isolated from a stone and sediment control sample were *C. albicans*, probably a human contamination and unidentified yeast respectively, and both of which did not produce beer.

Fifth – it is unlikely that these yeast cells originated from the soil or from handling contaminations. At least in the case of *Nakaseomyces delphensis,* the closest yeast species to the two isolated yeast strains EBEgT12, EBEgB8, was reported to be found on figs, and rarely in soil (81). Furthermore, the possibility that the source of the yeast cells is a contamination from modern beer is unlikely, since besides the *S. cerevisiae* yeast (TZPlpvs2), all the other isolated yeasts are not commonly used in the modern beer industry, and thus could not have derived from modern unfiltered beer. It should also be noted that although TZPlpvs2 is *S. cerevisiae,* its sequence is clearly different than commonly used *S. cerevisiae* laboratory strains, further excluding the possibility of contamination.

Taken together, we suggest that the evidence supports the authenticity of the yeast strains isolated from ancient vessels as ancient beverage yeast. We assume that the large amount of yeast cells that grew during a repeated series of fermentations in these vessels, in antiquity, were absorbed into the nano-pores of the vessels, altered the composition of microorganisms population, and remained as micro-colonies which continued to grow and survive over millennia in the ceramic matrix, based on occasional supply of moisture and nutrients.

In support of this assumption is the well-known fact that yeast in many traditional beer production methods, it is common to use the residues within vessels to serve as “starters” for the production of the next batch of fermented food. This technique is described in ancient inscriptions (57) and is still being used in modern traditional beer brewing techniques (22), and for the production of wine (82), yoghurt (52) and bread (67). Practically speaking, the ancient producers, using selection processes, domesticated yeast and bacteria that produced “good” and tasty fermented food in, similar to what was in the domestication of plants and animals were selected for. This perhaps could explain the findings that EBEgT12 and EBEgB8, isolated from Egyptian vessels from En-Besor, show high genetic similarities to each other, yet, they differ in several of the hallmark genes, typical of beer producing yeast strains, which seem to be mirrored in the quality of the beer that they produced. While EBEgT12 beer contained aromatic and flavor compounds, the beer made from yeast strain EBEgB8 had mildly spoiled aroma and flavors. Possibly, both of them were included in the original brew, complementing each other, or maybe, EBEgB8 represents the undomesticated ancestor of EBEgT12, before it was selected for “good” beer making as in the case of the domestication process of *S. cerevisiae* (66). These questions may be answered by additional isolation and analysis of yeast strains from more beverage containing vessels, which will shed further light on the yeast domestication processes.

In addition, the two yeast strains isolated from the Persian period vessels, RRPrTmd13 from the mead container and RRPrNerP7 from an oil lamp, show overall high similarity to each other, including similar genes associated to wine production (SI Appendix, Table S3). However, they diverge in growth under fermenting conditions, and in the quality of the beer they produced. In this case, we speculate that RRPrNerP7 represents the wild yeast ancestor, which naturally resides on olives (4, 29), while RRPrTmd13 is a domesticated descendant, which was selected for “successful” mead production. It might also suggest that wine and oil were prepared at proximal sites.

Regarding the red colony yeast TLVEgRD4 (*R. glutinis*), we suggest that it contaminated the ancient beverage, as happens today in modern traditional beers (20), or perhaps, although less likely, it was part of the beer sediments and contributed to its flavors.

In summary, based on all the above, we propose that it is highly likely that yeast strains EBEgT12, RRPrTmd13 and TZPlpvs2 are the descendants of the original ancient beverage producing yeast strains. We are less confident about TZPlpvs7, EBEgB8 and TLVEgRD4, which are perhaps descendants of contaminators of the ancient beverages. The yeast isolated from lamps originated, most probably, from yeast grew in the oil and the remaining yeast strains from sediments and stones, are probably wild yeasts.

It should be noted that for comparative reasons, the beers were brewed for these analyses using single yeast strains only, with a standard modern recipe. It is possible that brewing beverages using traditional recipes, ingredients and mixtures of the yeast strains and including those seemingly less fit for beverage production would have improved the brew quality. Moreover, it is highly likely that the yeast strains we isolated here represent only a portion of the rich variety of microorganisms that originally inhabited the vessels and contributed to the fermentation processes of the ancient beverages.

In conclusion, we show here that isolating, growing and studying fermenting microorganisms from ancient vessels in order to expand archaeological knowledge of ancient diet and food-related technologies, is feasible. These results, which allow a more precise recreation of ancient-like beverages than ever before, unlock enormous potential for the study of a broad range of food-related issues in antiquity. This includes expanding the knowledge about the ancient diet of diverse societies in many periods and locations, the study of the functions of ancient vessels, facilities and infrastructures, understanding links between cultures or identity groups and technological transfer between them, uncovering trade routes, food preparation technologies, and even insights on the actual somatic aspects (aroma and flavors) of ancient foods and beverages. Furthermore, the findings here might open new avenues in archeological research since we speculate that isolation of microorganisms from ancient remains is not limited to yeast only and it would be even easier to isolate bacteria due to their remarkable surviving abilities. Thus, this kind of approach can most probably be expanded to a broad range of topics, from disease-borne bacteria to food associated bacteria such as those used in fermented beverages, cheese and pickles.

The next steps of the research, currently conducted in our lab, will include “fingerprinting” of modern and ancient vessels which contained various kinds of fermented food and liquids. This is performed using combined microbiome-like DNA analysis and microorganism’s isolation which, we believe, will provide valuable data on the dating, identification, characterization of food containers and ingredients, and even the reconstruction of ancient diets.

## Materials and Methods

### Yeast growth

Unless otherwise mentioned, the yeast strains used in this work were routinely grown from a single colony either in liquid YPD medium (Difco, USA) at 30°C, under aerobic conditions with agitation (250–300 rpm) or on solid YPD medium containing 2% w/v Bacto-agar (Difco, USA) incubated at 30°C. Stocks of yeast strains were kept in −80° C in 50% glycerol.

### Preparation of control modern vessels for yeast isolation

A modern clay vessel was broken into equally sized and shaped pieces and divided into two groups; Group A pieces were buried “as is”, in three pits which were 30 cm deep and with a 2 meter space between them in a city garden. Group B shards were buried in the same way in a different city garden, situated several hundred meters away. Prior to covering them up, the pieces of group B were sprayed with 300ml of unpasteurized lager beer with a vital colony of the branded strain “Fermentis – WB-34/70”. After six weeks the pottery from both sites were retrieved and sent to the lab for yeast cell revival and isolation.

### Yeast isolation from vessels and control samples

The vessels were entirely flooded with rich YPD medium (Difco, USA) and incubated at room temperature for 7 days. Then, samples from the medium were streaked on selective agar plates for fungal isolation (NOVAmed BA-114, Israel) and incubated at 30 °c for 12 to 48 hours. Yeast colonies growing on the plates were re-plated on solid YPD agar plates, containing 2% w/v Bacto-agar (Difco, USA). Colonies were picked for further analysis.

### Electron microscopy

Ceramic samples were cut using a diamond disc power cutter (Dremel). The surface morphology of the archeological ceramic samples was examined using the FEI Quanta 200 scanning electron microscope situated in the core facility of the Hebrew University Medical School in Ein Kerem. Samples were first sputtered by Au/Pd (SC7620, Quorum Technologies). Images were then taken with a secondary electron detector at 10k – 40k × magnification using a 10-30 kV accelerating voltage and an objective lens aperture of 30-20 μm.

### DNA purification

Yeast cell DNA isolation was performed as previously described (32). Briefly, 10ml of overnight cultures were centrifuged at 3000 rpm for 5 minutes and washed in sterile water. The cells were treated with 200µL of phenol chloroform, 0.3g of acid washed glass beads and 200µL of Smash and Grab solution (32), and lysed using a vortex for 3 minutes. after which TE buffer was added. The cells were centrifuged, and the aqueous layer containing the DNA was transferred to 1ml ethanol, washed and suspended in TE buffer. 1µL of RNase (10mg/ml DNase- and Protease-free RNase, ThermoFisher Scientific) was added, and the solution was incubated at 37°C for 5 minutes. 10µL of ammonium acetate (4M) and 1ml of ethanol were then added, and the solution was washed, and suspended in 100µL of TE buffer. The extracted DNA was stored at −20°C. DNA quantification was carried out on a Synergy H1 microplate reader (BioTek Instruments, Inc., Vermont, USA), using a Take3 micro-volume plate.

### Internal Transcribed Spacer (ITS) analysis

The ITS region of the yeast was amplified using standard Illumina primers as described in the Earth Microbiome Project web site (http://www.earthmicrobiome.org/protocols-and-standards/its/). PCR fragments were Sanger sequenced by the inter-departmental sequencing unit of the Hebrew University. The sequences were identified by BLAST analysis against the ISHAM barcoding database (http://its.mycologylab.org) and the NCBI database (https://blast.ncbi.nlm.nih.gov/Blast.cgi).

### DNA Sequencing

Sequencing was performed in the inter-departmental unit at the Hebrew University, Hadassah Campus. Libraries were prepared by using a Nextera XT DNA kit (Illumina, San Diego, CA), and DNA was amplified by a limited-cycle PCR and purified using AMPure XP beads. The DNA libraries were normalized, pooled, and tagged in a common flow cell at 2×250 base-paired-end reads using the NextSeq platform.

### Genome assembly

Raw reads are available on Genbank (PRJNA449847). Illumina adaptors were removed with Trimmomatic 0.36 (6). The quality of the reads was determined using FastQC (http://www.bioinformatics.bbsrc.ac.uk/projects/fastqc). *De-novo* assembly was then carried out with the Celera assembler 8.3rc2 (62) and non-target-species scaffolds were excluded using BlobTools V1 (45). The sequencing and genome assembly effort was targeted at obtaining assemblies contiguous enough to derive protein coding gene data for phylo-genomic analyses (SI Appendix, Table S5). The resulting genome assemblies had coverages of 36-240X, and N50 values of 2842-15623 bp, SI Appendix, Table S5). These data provided us with at least 2702 protein coding genes per sample with at least 294 AA median length. The gene count variation could be both the result of ploidy differences or of genome assembly artifacts, and may cause the underestimation of one to one orthologs. However, we were still able to curate a large and high quality one to one ortholog gene subset to perform the phylogenomic analyses.

### Beer preparation

For beer production comparison we followed a common standard recipe (44) where only the yeast strain was changed. Water (5L) was heated to a pasteurization temperature of 72° C. Malt extract was added to a final concentration of 100gr/L, while thoroughly stirring, and allowed to infuse together for 30 min in temperatures between 63-67° C. The solution was then heated to 100° C, and once boiling has occurred 1gr/L of hops were added. The mixture was allowed to boil for 45 more minutes, followed by the addition of another 1gr/L of hops. The mixture was then heated for an additional minute. Previously prepared ice-cold water was then added to the mixture, and the prepared wort was transferred to a sanitized fermenter and brought to a final volume of 10L. The wort was left at room temperature for 30 min, and then overnight cultures of yeast were added. Fermentation typically began within 12-48h, and the mixture was left untouched for a week.

### HS-SPME Procedure and GC-MS analysis of beer

The method we used was based on the method described by Rodriguez et al. (70). Beer bottles were cooled at 4°C to prevent loss of volatiles. Beer sample (6 ml), a magnetic stirrer, 100 μL of an internal standard (5ppm 2-Octanol) and 1.8 g of NaCl were added to 20-mL SPME headspace vials and were sealed with PTFE/Silicon septum (Supelco). The samples were then incubated for 10 min at 44.8 °C in a water bath on a heating plate and stirred by magnetic stirrer. The septum covering the vial headspace was pierced with the needle containing the SPME fiber, retracted and the fiber was subsequently exposed to the headspace for 47 min at 44°C, then inserted directly into the GC-MS injection port. SPME Fiber 50/30-μm Divinylbenzene/ Carboxen/ Polydimethylsiloxane (DVB/CAR/PDMS) 2-cm length and manual holder were purchased from Supelco (Sigma Aldrich). The analyses were performed using a gas chromatographer (Agilent 6890N) fitted with a splitless injection with a liner suitable for SPME analysis and an Agilent 5973 mass spectrometer (MS) detector in Full-Scan mode. An Agilent MSD ChemStation Software was used to control the gas chromatograph (G1701-90057). Ultra-high purity grade helium was used as the carrier gas at a flow rate of 1 mL/ min. Samples were analyzed on a DB-5MS UI column (30-m×0.250-mm i.d.×0.25-μm film thickness) from Agilent. The oven temperature was programmed as follows: 40 °C as initial temperature, held for 5 min, followed by a ramp of temperature at 4 °C/min to 60 °C and then at 8 °C/min to 200 °C held for 15 min; holding at this temperature for 5 min. An electron impact ionization technique was used at 70 eV. The detector range of scan was from m/z 10 to 250. Suggestions for the identification of the detected peaks were carried out by Wiley mass spectrometry database. Peak areas were calculated using the integration order in the ChemStation Software. For each sample, we determined the peak area for 2-octanol standard and ethanol, as well as for 35 aroma compounds usually found in beer. Following the integration of the 2-octanol peaks we were not satisfied with its repeatability between technical repeats. Thus, we decided to use the peak areas of ethanol, which was separated clearly and was highly correlated to its determination by distillation in our lab, as internal standard for each sample. To achieve normalized ethanol peak areas, we divided the peak area of ethanol by its concentration (%) determined by distillation, for each beer sample. Finally, the relative peak area for each compound was calculated by dividing the peak area of the compound to that of the normalized ethanol peak area, and multiplied in 1000, to get more presentable numbers. This allows us a presentation of qualitative analysis of relative peak areas for each compound across our samples. The results presented, and the statistical analysis were done by averaging the three biological samples for each yeast strain.

### Determination of carbohydrates in beers

Stock solution of Phenol (J&K Scientific Gmbh), 0.05g/mL and D(+)-Glucose (Merck), 100μg/mL were prepared. Glucose Standards-Aliquots of 1mL, 2mL, 3ml, 4mL, 5mL, 8mL, 10mL 13mL and 20mL of the glucose stock solution were pipetted and transferred into nine 30mL beakers. Adequate amount of distilled water was added to make a final volume of 20mL. Each solution (2mL) was measured and transferred into 10 test tubes. The phenol (2mL) and 10mL of the concentrated 95%-97% sulfuric acid (Merck) were pipetted and added to each of the 10 test tubes. A light orange color developed and the tube was allowed to stand for 10mins. The solutions were then transferred into 1-cm path length cuvettes and the absorbance’s were measured at 485nm with a UV spectrophotometer (Genesys 10S UV Vis, Thermo). For measurements, one mL of beer was measured and transferred into 1L volumetric flask. Distilled water was added to make 1000mL solution. Aliquots (2mL) were transferred into test tubes and mixed with 2mL phenol solution and 10mL concentrated sulfuric acid. A light orange color developed, and the absorbance was measured at 485nm after 10 mins. Results were determined by averaging triplicate measurements. Ethanol concentration in beers samples was determined using a digital distillator Super Dee and an electronic hydrostatic balance model Super Alcomat (Gibertini, Italia). pH values of beer samples were measured using a pH meter Hana HI 2211 (Hanna Ins.).

To analyze their spectrophotometric properties the beer samples were degassed and centrifuged followed by a spectrophotometeric (Genesys 10S UV VIS Thermoscientific) measurement at 430 nm (quartz cuvettes 10 mm). Beer color was calculated by two scales SRM and EBC (SRM=Absorbance*12.7; EBC= Absorbance*25.0). To determine the beers’ density, we used a hydrometer (“Alla” Franc).

### Beer tasting

The flavor and aroma assessments were performed according to the BJCP’s judge procedure manual (https://www.bjcp.org/judgeprocman.php) as following: A 100ml sample was served to the assessors in identical vessels to prevent variations of aroma and flavor compounds distribution. The assessors then recorded their impressions discreetly on a recognized form, to avoid bias between the tasters. The forms, inter-divided to the subjects of: appearance, aroma, flavor and overall impression, were then collected, summarized and processed. The summary ignored the appearance and overall impression sections, as well as hop flavor and aroma entries and focused primarily on known fermentation byproducts and sugar residue compounds. All “named entries” on the forms (-such as Caramel/Fruity/etc.) come with a notation of the strength of the flavor/aroma derives from on a scale of 1-5 (left column on the evaluation form) and averaged by 5 – testers.

### Statistical analysis

Statistical analysis was performed using R (https://www.r-project.org) and Prism Graphpad 7 (https://www.graphpad.com/scientific-software/prism/). Differences between growth curve in wort medium (Fig 3) were calculated using R “growthcurver” package (https://cran.r-project.org/web/packages/growthcurver/vignettes/Growthcurver-vignette.html) by fitting the growth data to the logistic equation. The r parameters of each curve were compared either to that of SafAle S.04, a modern beer yeast which served as control, or to each other using Principal Components Analysis (PCA, R prcomp() command). For significance distances from the control growth curve we used the Student’s t-test. Differences between aromatic and flavor compounds in beer produced by the isolated yeast strains (Fig 4) and aromas and flavors of these beer were compared by clustering analysis using R function hclust()with method “complete” and dist() function with method “euclidean”. The dendograms and clusters were created using Ward hierarchical clustering with bootstrapped *P-values* using R pvclust() method from R package pvclust, with parameters: hclust=“ward.D2” and method.dist=“Euclidean.

## Acknowledgments

We thank the Israel Antiquities Authority and the Israel Nature and Parks Authority for assistance, and the staff and team members of the Tell es-Safi/Gath Archaeological Project (Bar-Ilan University) and the Ramat Rachel Archaeological Project (Tel Aviv University). We also thank the members of the Beer Judge Certification Program (BJCP) Omer Basha, Noam Shalev, Ephraim Greenblat and Roi Krispin and the beer experts and brewers Shmuel Naky, Eyal Grossman and Yisrael Atlow for tasting and evaluating our beers. We thank the team of the Interdepartmental Core Unit of the Hebrew University at the Hadassah Ein Karem campus; Edi Berennshtein, for assistance with the electron microscopy and Abed Nasereddin and Idit Shiff for NGS sequencing. Thanks also to Marina Faerman, Philipp Stockhammer and Christina Warinner for critical reading of the manuscript and to Ayelet El-Boher, Sharon Ben-Hur and Maayan Margulis for their technical assistance. Last, we thank Kedma winery, Kfar Uria, Israel for supplying fragments of a wine clay container that was unused for the last two years.

## Authors contributions

RH and MK designed the experiments. TA, DG, SCG, ER, MS and RK isolated and characterized the yeast strains. AS^1^ analyzed the genomic data. IG produced the beer. YP, AMM, YG and OL contributed the vessels from their excavations and shared their archeological insights and AS^2^ participated in archeological samples preparations. ED and AP analyzed the beer compounds. TBG performed the electron microscopy imaging. RH, MK and SCG analyzed the results. RH, MK, YP, AMM, YG, AS and ED wrote the manuscript. RH, AMN and TBG prepared the figures.

**The authors declare no conflict of interest.**

## Data deposition

Genetic sequence analysis input and output files are available in the FigShare repository (https://figshare.com), with DOI numbers as provided in the following legends.

## Comments

1. *Nakaseomyces delphensis* has the following synonyms: *Saccharomyces delphensis, Dekkeromyces delphensis, Guilliermondella delphensis, Kluyveromyces delphensis, Zygofabospora delphensis*(*3, 10*) (http://www.mycobank.org/name/Nakaseomyces%20delphensis).
2. *Hyphopichia burtonii* has the following synonyms: *Pichia burtonii,, Pichia burtonii, Endomycopsis burtonii, Boidin, Candida armeniaca-cornusmas, Candida fibrae Nakase, Cladosporium fermentans, Sporotrichum anglicum, Sporotrichum carougeaui, Trichosporon behrendii, Trichosporon beijingense, (*http://www.mycobank.org/Biolomics.aspx?Table=Mycobank&Rec=36231&Fields=All)
3. ***SSU-rRNA***: The patristic distance (substitutions per base) between the isolate and its closest relative, given with the probability that this is an intraspecific tree distance.
4. **Phylogenomic tree**: The phylogenetic position of the isolate in the phylogenomic tree, provided with the branch support of this relationship. Node supports are bootstrap percentages.
5. **Whole genome blast**: The closest blast match of most contigs and the proportion of contigs that are assigned to this match. See supplementary material for description of percent identities of closest matches.
6. **Node support (Bootstrap Percentage, BP):** The percentage of bootstrap tree that had the same topology as the maximum likelihood tree for a given node.
7. **Scaffold prop:** The proportion of scaffolds that had the taxon as their first blast hit.

